# EEG-based Assessment of Long-Term Vigilance and Lapses of Attention using a User-Centered Frequency-Tagging Approach

**DOI:** 10.1101/2024.12.12.628208

**Authors:** S. Ladouce, J. J. Torre Tresols, K. Le Goff, F. Dehais

## Abstract

**Objective:** Sustaining vigilance over extended periods is crucial for many critical operations but remains challenging due to the cognitive resources required. Fatigue and other factors contribute to fluctuations in vigilance, causing attentional focus to drift from task-relevant information. Such lapses of attention, common in prolonged tasks, lead to decreased performance and missed critical information, with potentially serious consequences. Identifying physiological markers that predict inattention is key to developing preventive strategies.

**Approach:** Previous research has established electroencephalography (EEG) responses to periodic visual stimuli, known as steady-state visual evoked potentials (SSVEP), as sensitive markers of attention. In this study, we evaluated a minimally intrusive SSVEP-based approach for tracking vigilance in healthy participants (N = 16) during two sessions of a 45-minute sustained visual attention task (Mackworth’s clock task). A 14 Hz frequency-tagging flicker was either superimposed on the task or absent.

**Main results:** Results revealed that SSVEP responses were lower prior to lapses of attention, while other spectral EEG markers, such as frontal theta and parietal alpha activity, did not reliably distinguish between detected and missed attention probes. Importantly, the flicker did not affect task performance or participant experience.

**Significance:** This non-intrusive frequency-tagging method provides a continuous measure of vigilance, effectively detecting attention lapses in prolonged tasks. It holds promise for integration into passive brain-computer interfaces, offering a practical solution for real-time vigilance monitoring in high-stakes settings like air traffic control or driving.

**Highlights:**

- Fluctuations in the SSVEP response to a continuous frequency-tagging flicker presented throughout a 45-minutes long vigilance task was used to characterize lapses of attention
- The frequency-tagging flicker superimposed to the task was designed to be minimally intrusive thanks to its low luminance and low contrast
- Neither the user experience nor the task performance were altered by the presence of the frequency-tagging flicker
- The SSVEP response outperformed measures of individual alpha peak and theta band activity in the distinction of attentional lapses (missed events) from successful target event identification
- The present findings have implications for the design of Human-Computer Interfaces aimed at monitoring (in)attention during the performance of monotonous routines (e.g., surveillance, driving,…)

## 1. Introduction

Vigilance, defined as the capacity to sustain attention over prolonged periods (Oken et al 2006; Mehrabi and Kim 2022), plays a critical role in high-stakes environments where timely responses to new information are essential (Mehta and Parasuraman 2013; Small et al 2014; Lin et al 2014; Zheng et al 2011). However, vigilance is not a static state; it fluctuates due to various factors and typically declines over the course of extended tasks, often resulting in lapses of attention (Hancock, 2017). These lapses, characterized by reduced responsiveness or failure to detect critical information, can lead to serious real-world consequences, such as impaired driving performance (Papadelis et al 2007).

Two main hypotheses have been proposed to explain lapses in attention. The first posits that lapses arise from the depletion of top-down executive control resources necessary to maintain focus over time (Warm et al 2008; Kamzanova et al 2014; Thomson et al 2015). As these cognitive resources diminish, sustaining attention becomes increasingly effortful. The second hypothesis suggests that lapses occur due to task monotony, which leads attention to drift away from task-relevant information and fosters a “mindless” operational mode (Manly et al 1999). Factors such as habituation to repetitive stimuli (Möckel et al 2015), reduced motivation (Brosowsky et al 2023), and increased boredom (Esposito et al 2022; Esterman et al 2016) collectively contribute to this progressive task disengagement.

In line with these frameworks, vigilance fluctuations and lapses in attention have been linked to cognitive fatigue (Tian et al 2018; Trejo et al 2015; Tran et al 2020), mental workload (Borghini et al 2014), mind-blanking (Ward and Wegner 2013), mind-wandering (Chun et al 2011; Groot et al 2021; Andrillon et al 2021), and distractions from task-irrelevant stimuli (Pershin et al 2023). Furthermore, chronobiological factors such as circadian rhythms have been implicated in these fluctuations (Czisch et al 2012; Gibbings et al 2021; Hudson et al 2020; Jagannathan et al 2018; Xie et al 2023). Regardless of the terminologies or theoretical frameworks employed, identifying predictive markers for vigilance lapses remains a critical goal for advancing neuroassistive technologies aimed at enhancing safety and performance in attention-demanding tasks.

Various sustained attention tasks have been employed to study the neural correlates of vigilance decrement, frequently revealing time-on-task effects such as increased error rates and slower reaction times (Manly et al 1999; Robertson et al 1997; Drummond et al 2005; Mackworth, 1948). For instance, the Sustained Attention to Response Task (SART, Torkamani-Azar et al 2020), combined with experience sampling, has demonstrated that task-unrelated thoughts are related to increases in alpha, theta, and delta power, as well as an imbalance between the default mode network and its anticorrelated network (Groot et al 2021). Similarly, Molina et al (2019) examined EEG dynamics during a 45-minute Psychomotor Vigilance Task (PVT, Dinges and Powell 1985; Drummond et al 2005), finding that increases in alpha, theta, and beta power were associated with slower response times, reflecting a decline of task performance.

In another EEG study, Esposito et al (2022) used Mackworth’s Clock Task (MCT, Mackworth, 1948) to investigate vigilance decrements and found that frontal theta power positively correlated with reaction time, while parietal theta power was negatively correlated with detection rates, underscoring distinct neural correlates of vigilance lapses. Boksem et al (2005) similarly recorded EEG data during a three-hour visual attention task, finding that subjective fatigue ratings and power in theta and lower-alpha bands increased significantly over time, reflecting fatigue accumulation. Additionally, in a four-hour EEG study, Wascher et al (2014) observed that mental fatigue manifested through a progressive rise in alpha power within the first hour, followed by sustained increases in frontal theta activity. These increases in theta reliably reflected cognitive fatigue and changes in attentional engagement, indicating that vigilance decrements could be mapped through spectral power dynamics.

Furthermore, specific neurophysiological markers have been identified as predictors of impending lapses in vigilance. For example, O’Connell et al (2009) reported increased alpha power in the right inferior parietal cortex up to 20 seconds before errors in a continuous temporal expectancy task, suggesting it as a neural marker for approaching lapses. Using a prolonged Flanker task, Eichele et al (2010) documented a gradual decline in N2 amplitude starting several trials before errors. Similarly, Shou et al (2015) found that pre-stimulus alpha activity predicted errors in a prolonged color-word matching Stroop task. In a notable study involving Mackworth’s Clock Task, Martel et al (2014) observed distinctive neural patterns preceding lapses, including increased alpha power approximately 10 seconds before a missed target, likely reflecting a shift toward an internally focused attentional state. The authors also reported a reduction in the P3 component, an indicator of attentional resource allocation, in response to prior events occurring up to five seconds before lapses, suggesting a progressive disengagement from the task.

While these findings offer valuable insights into the neural mechanisms underlying attention fluctuations, they face practical challenges for real-world monitoring Dehais et al (2020). Although various EEG frequency indices have been linked to mind-wandering and performance decrements, they lack the specificity needed to reliably predict attentional lapses. This limitation arises because frequency bands such as alpha, theta, and beta are associated with a broad range of cognitive phenomena (Kahana, 2006), complicating efforts to isolate attentional lapses based solely on these measures. Additionally, studies suggest that these spectral markers often lack temporal stability (Roy et al 2013, 2016), which may confound electrophysiological responses with time-on-task effects. Moreover, extracting precise, time-sensitive features like ERPs requires knowledge of specific event timings, posing significant challenges for real-time applications.

Steady-State Visual Evoked Potentials (SSVEPs) refer to sustained oscillatory responses in the visual cortex that match the frequency of a flickering stimulus. In paradigms where multiple stimuli flicker at distinct frequencies are presented, each flicker elicits a separate “frequency-tagged” SSVEP, allowing researchers to track attention to each stimulus independently (Norcia et al 2015). Early work by Morgan et al (1996) showed that directing attention toward a specific flicker leads to a substantial increase in SSVEP amplitude, underscoring the sensitivity of SSVEPs to spatial selective attention. In a follow-up EEG–fMRI study, this attention-related enhancement of the SSVEP response was localized to ventral and lateral extrastriate areas in the occipital lobe (Hillyard et al 1997). Further studies found that both the amplitude and latency of SSVEPs depend on flicker frequency (Müller et al 1998; Ding et al 2006), luminance–color contrast (Russo and Spinelli 1999), and task demands (e.g., increased amplitude during visual memory encoding; Peterson et al 2014). In addition, individual differences in alpha peak frequency (Gulbinaite et al 2017) and distinctions between alpha- and gamma-band responses (Gulbinaite 2019) reflect varying resonance properties across participants and frequencies. Keitel et al (2017) extended these findings showing that the visual system systematically phase-locks to quasi-rhythmic stimulation across theta, alpha, and beta bands, with spatial attention further enhancing synchronization at lower frequencies. These findings thereby emphasized the continuous, rather than strictly rhythmic, nature of the neural entrainment in early visual cortices reflected by SSVEP. In a recent large-scale reanalysis of eight studies (N = 135), Adamian and Andersen (2024) demonstrated that attention consistently led to an increase in SSVEP amplitude. By providing a continuous and sustained measure of visual attention, SSVEPs emerge as a promising marker for real-time monitoring of vigilance fluctuations.

To the best of the authors’ knowledge, only two studies have specifically examined the role of SSVEPs in vigilance assessment. Silberstein et al (1990) investigated the topographic distribution of SSVEPs using a 13 Hz flicker during a visual vigilance task, in which participants monitored a screen for rare target stimuli. Their key finding was that during periods of heightened attention, such as when participants anticipated the appearance of a target, SSVEP amplitude significantly decreased, particularly in the occipito-parietal and right prefrontal regions. This decrease was most pronounced approximately 10 seconds after stimulus onset and persisted for up to 50 seconds. However, the relatively short task duration (nine minutes split in three trials of three minutes each) may not have been sufficient to induce attentional lapses (Möckel et al 2015; Lim and Kwok 2016). Moreover, since no behavioral data on lapses were recorded, a direct relationship between changes in SSVEP amplitude and lapses of attention could not be established.

More recently, O’Connell et al (2009) examined the temporal dynamics of brain activity preceding lapses of attention during a continuous temporal expectancy task. In this task, participants monitored a stream of alternating patterned stimuli flickering at 25 Hz to detect infrequent target stimuli that were 40% longer in duration than the frequent stimuli. Although changes in other electrophysiological markers, such as increased alpha activity (8–14 Hz) and reduced frontal P3, were observed to precede attentional lapses (starting 20 seconds before a missed target), variations in SSVEP amplitude were not associated with these lapses. SSVEP dynamics were computed from consecutive two-second segments and further smoothed over time, which may have reduced the sensitivity of SSVEP features. This reduced sensitivity could potentially explain the discrepancy with earlier findings that reported differences in SSVEP amplitude preceding errors and correct trials.

While these studies provide relevant insights, they do not offer conclusive evidence for using SSVEP to track attention fluctuations or predict attentional lapses. Beyond certain limitations in experimental design, one reason for the absence of significant results may be that the flickering of the stimuli was too pronounced. Prolonged exposure to flickering stimuli can negatively impact user experience due to their distracting and visually intrusive nature (Zemon and Gordon, 2006; Wu and Lakany 2013; Duszyk et al 2014; Ladouce et al 2021). High contrast and luminance of flickering elements tend to capture visual attention, disrupting task-focused visual exploration strategies (Kinchla and Wolfe 1979), and leading to issues such as eye strain (Zhu et al 2010), mental fatigue (Makri et al 2015), and increased drowsiness (Cao et al 2014; Ortner et al 2011; Patterson Gentile and Aguirre 2020). These issues could distract participants from the task, introduce bias in their attentional state, and adversely affect vigilance.

In order to circumvent these issues and enable the use of SSVEP to tag fluctuations in attention and detect attentional lapses in a non-intrusive way, two main strategies have emerged: Rapid Invisible Frequency Tagging (RIFT) and amplitude modulation depth reduction. RIFT capitalizes on high-frequency flickers to elicit neural responses to stimuli that are imperceptible to the user, showing promise for investigating cognitive processes (Zhigalov et al 2019; Drijvers et al 2021; Pan et al 2021). However, this approach requires specialized high-refresh-rate projectors and high-density neuroimaging systems, imposing significant technical constraints and limiting broader application (Arora et al 2024). In contrast, amplitude modulation depth reduction focuses on decreasing the contrast and intensity of flickers by lowering their amplitude modulation depth (Mouli and Palaniappan 2016). Recent research indicates that flickers of lower contrast can achieve higher classification performance in SSVEP-based Brain-Computer Interface (BCI) applications while maintaining user comfort (Ladouce et al 2022, 2021; Cabrera-Castillos et al 2023). Furthermore, amplitude modulation depth reduction at the threshold of conscious perception can effectively frequency-tag spatial attention with high precision (Ladouce and Dehais 2024).

The present study aims to build on the foundation of SSVEP research by applying a minimally intrusive 14 Hz frequency-tagging flicker to monitor vigilance lapses during a 45-minute sustained attention task. We employed Mackworth’s Clock Task (Mackworth, 1948) as a continuous visual attention paradigm, simulating prolonged monitoring scenarios. Unlike previous studies that have relied on extended Fourier-transformed EEG segments, potentially limiting the temporal resolution required for precise vigilance assessment, our approach harnesses advanced signal processing and feature extraction techniques. This enables us to capture moment-to-moment modulations in SSVEP responses, facilitating a continuous and detailed assessment of vigilance lapses. In addition to behavioral data, we will collect subjective reports to assess the impact of the flicker on user experience. We expect that the refined SSVEP approach will offer a more continuous and accurate measure of vigilance compared to traditional methods, and our SNR-based analysis will be benchmarked against classical alpha and theta band measures, which have both been reported as indicators of vigilance fluctuations. We hypothesize that the signal-to-noise ratio (SNR) of the SSVEP response recorded over occipital electrodes will predict lapses of attention as indicated by behavioral performance measures (misses vs. hits).

## 2. Methods

### 2.1. Participants

Sixteen participants (mean age = 27.5 years, SD = 5; 14 right-handed; 11 males) took part in this study. The sample size was determined based on prior research investigating SSVEP responses in similar experimental contexts. Silberstein et al (1990) used a sample of 15 participants for initial investigations of long-term SSVEP fluctuations, while subsequent studies employed sample sizes between 15 and 21 participants to explore continuous SSVEP dynamics and other EEG markers of attention lapses (Silberstein et al 2000; O’Connell et al 2009; Esposito et al 2022). Participants received a 20€ voucher as compensation for their time. Written informed consent was obtained from all participants before the experiment, and they were informed of their right to withdraw at any time. Exclusion criteria included a history of epileptic seizures, visually-induced migraines, and general photosensitivity, which were screened during recruitment to ensure participant safety due to the flickering stimuli used in the study. All participants had normal or corrected-to-normal vision. All participants were healthy individuals without any cognitive or motor impairments and reported no sleeping disorders. Nine participants had prior experience with BCI and EEG recording protocols. The study was approved by the Ethics Committee of the University of Toulouse (CER approval number 2023-749) and conducted in accordance with the principles embodied in the Declaration of Helsinki.

### 2.2. Paradigm

The well-established Mackworth Clock Task (MCT, Mackworth (1948)), renowned for its robustness in capturing behavioral markers of vigilance fluctuations and lapses of attention, was used in this study. Participants were seated comfortably in front of a monitor and instructed to maintain focus throughout the 45-minute task. The task featured a clock-like circular array of 96 linearly spaced positions displayed consistently on the screen, with a moving indicator (a red dot) that advanced in discrete steps every 800 ms (Figure 1).

**Figure 1:**
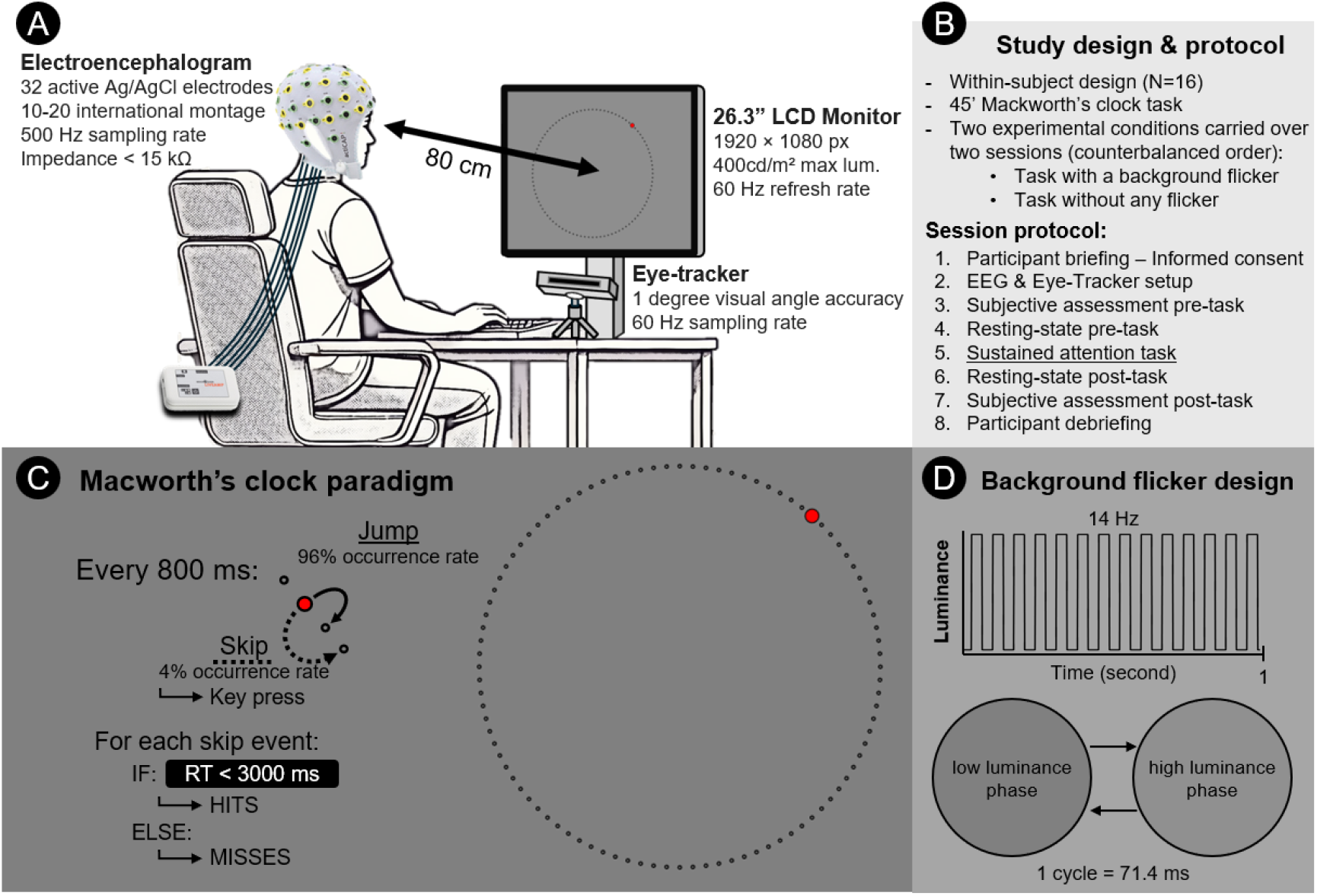
A. Experimental setup: Participants were seated comfortably in front of a 26.3” LCD monitor where the experimental task was presented. They were equipped with 32 electrodes for EEG recording, and their gaze was tracked using a head-mounted eye-tracker. B. Study design and protocol: Information regarding the design and procedure of an experimental session. C. Experimental paradigm: The Mackworth Clock Task was used to investigate vigilance fluctuations and lapses of attention in a long-term sustained attention context. Participants monitored a red dot moving clockwise around a 96-position clock. When the dot skipped a position (double jump or skip), participants were instructed to press a button within five seconds (hit trials); otherwise, the trial was considered a miss. The occurrence rate of skip events was set at 4%. D. Flicker design: The transparent flicker was superimposed on the task, modulating the luminance of the entire screen at a rate of 14 Hz using a square wave function. The low and high luminance phases are shown below.

Participants were required to monitor the movement of this indicator and respond as quickly and accurately as possible when the indicator skipped a step, simulating an unexpected or irregular movement. This target event could appear at any position on the circular array and followed a set of pseudo randomization rules. Specifically, the list of all 3240 steps covering the whole 45 minutes duration of the task was divided into 18 series of 180 events, within which target jumps had an occurrence rate of 4%. Target events were not allowed within the first four steps (or 3.2 seconds) of a series, and the presentation of consecutive targets were separated by at least eight regular steps. The target occurrence rate and task duration were based on previous studies that effectively induced time-on-task effects using the MCT paradigm Esposito et al (2022).

Participants were instructed to press a designated response button (spacebar) as soon as they detected a target event. Reaction times (RTs) to target events were recorded, and if no response was made within three seconds of a target event, it was registered as a miss. Responses made without a corresponding target event were registered as false alarms, although the low incidence of false alarms in most datasets precluded further analysis of this measure. Prior to the main experimental session, participants completed a 2-minute training session to ensure understanding of the task.

The aforementioned experimental protocol was repeated in two sessions: once with flicker and once without flicker, with the session order counterbalanced across participants. The duration of each session did not exceed 90 minutes. Each session included participant briefing, EEG setup, a first resting-state recording, baseline questionnaires, completion of the 45-minute Mackworth’s clock task, post-experiment questionnaires, a post-experiment resting-state recording, and a debriefing.

### 2.3. Frequency-tagging flicker design

A flickering visual stimulus (or frequency-tagging probe) was presented continuously throughout the task, covering the entire screen (LCD, 26.3 inches, 1920 × 1080 pixels, maximal luminance: 400cd/m2, 120 Hz refresh rate). The flicker frequency was set to 14 Hz, based on prior studies (Ladouce et al 2022, 2021), which found that flickering rates between 10 and 20 Hz reliably evoke SSVEP responses with a high signal-to-noise ratio (SNR). The choice of 14 Hz was also intended to avoid overlap with frequency bands commonly linked to sustained attention, such as theta, alpha, and gamma bands Clayton et al (2015). Specifically, the 14 Hz flicker was chosen to minimize interference with the alpha band (8-12 Hz), reducing potential confounds when comparing SSVEP with ongoing alpha-band activity. Since the analysis focused on the characterization of the spectral response at the fundamental stimulation frequency, a square-shaped waveform was preferred (Teng et al, 2011; Chen et al, 2019). The flicker was implemented using the FlickersOnTop (v1.0) software [https://github.com/neuroergoISAE/FlickersOnTop], which was chosen for its ability to overlay flickering stimuli seamlessly on top of other software windows, ensuring precise control over the flicker’s presentation without interrupting the experimental task. In the low luminance phase (flicker completely transparent), the task environment was composed of a grey background (value of 130 on the gray scale) whose luminance was 124cd/m2 (measured using a digital light meter from Extech Instruments). During the high luminance phase of the flicker period, the slightly brighter (155 on the gray scale) screen-wide flicker was presented on top of the window running the experimental task with 80% transparency. As such, the luminance of the grey background alternated between 124 and 146cd/m2 across the phases of the transparent flicker period (see Figure 1.D). The decision to use a relatively low contrast level was made to minimize potential eye strain and ensure participant comfort during prolonged exposure, while still maintaining a detectable SSVEP response^1^.This decision was informed by our previous work on optimizing user experience and SSVEP signal for BCI applications (Ladouce et al, 2021, 2022).

### 2.4. EEG data acquisition and processing

The EEG data were acquired using 32 active (Ag/AgCl) electrodes fitted according to the 10-20 international montage in an elastic cap, connected to a LiveAmp amplifier (Brain Products, Munich, Germany). All electrodes were referenced to the FCz electrode, with the Fpz electrode used as ground. The impedance of all electrodes was brought below 15 kΩ using conductive gel. EEG data were recorded at a sampling rate of 500 Hz with an online digital band-pass filter ranging from 0.1 to 250 Hz. At the onset of every experimental event (frequent jumps, infrequent double jumps), an event trigger was generated by the stimulus presentation program (MATLAB code available) and synchronized to the EEG data stream through the Lab Streaming Layer (LSL, Kothe et al 2024) data synchronization framework.

After data acquisition, the following signal processing pipeline was applied to all EEG datasets. First, the raw continuous EEG data underwent an offline 1 to 40 Hz bandpass filtering (zero phase, acausal, filter order: 1651, -6 dB cut-off frequencies at 0.5 and 40.5 Hz). The data were then re-referenced to the average of all channels. An infomax Independent Component Analysis (ICA, Makeig et al 1995) was then performed on the continuous data. To remain as close to the original data and mimic an online assessment scenario, no artifact cleaning was performed prior to ICA decomposition. The number of Independent Components (ICs) to compute was adjusted to match data rank deficiency stemming from the average referencing (Delorme and Makeig 2023). Next, we used the DIPFIT plugin in EEGLAB to compute an equivalent dipole model for each IC. Electrode locations were warped to fit a boundary element head model based on the MNI brain (Montreal Neurological Institute, QC, Canada). The dipole model also provides estimates of how much variance in each IC’s topography remained unexplained by the model, referred to as the residual variance (RV). Because EEG signals originating from cortical sources tend to exhibit dipolar patterns (Delorme et al., 2012), the RV measure has been used as an indirect index of ICA decomposition quality in technical reports (Klug et al., 2024). Thus, ICs with high RV (e.g., >20%) are less likely to represent genuine brain sources. We further identified artifactual ICs based on classification confidence scores from the ICLabel algorithm (Pion-Tonachini et al., 2019). Specifically, ICs were discarded if their confidence scores exceeded 80% in any of the following categories: ocular, muscular, heart rate, line noise, electrode, or other artifact. As a result of this pruning, a mean of 13.4 (SD = 2.45) ICs were discarded, resulting in an average of 18.6 (58%) remaining components per dataset. This artifactual IC pruning strategy is relatively conservative compared to the guidelines regarding the ratio of bad ICs proposed in (Klug and Gramann 2021). Among the remaining ICs, the residual variance (mean = 12.48%; range = 3.01–16.99%) indicates that these components plausibly reflect brain-generated signals.

### 2.5. Measures

#### 2.5.1. User experience assessment measures

Several subjective experience measures were collected both before and after the task to evaluate whether the flicker influenced sleepiness, eye strain, general fatigue, and perceived task difficulty. For measures that violated the normality assumption (tested using the Shapiro-Wilk test), a Wilcoxon signed-rank test was employed instead of a paired-sample Student’s t-test.

**-** **Karolinska Sleepiness Scale (KSS)**, Akerstedt and Gillberg (1990) was used to assess sleepiness, consisting of a single item measuring participants’ subjective alertness/sleepiness on a 9-point Likert scale. Sleepiness induced by the task was indexed by the difference between pre- and post-task scores.
**-** **Digital Eye Strain Questionnaire (DESQ)**, Mylona et al (2022) was administered before and after the task in both sessions. This binary (yes/no) screening questionnaire evaluates eye discomfort and vision problems related to prolonged screen exposure. The DESQ includes 12 items (e.g., blurry vision, watery eyes, headaches, sensitivity to light), with each reported symptom adding one point to the scale. The difference in the total symptom scores pre- and post-task was compared between conditions.
**-** **Visual Analog Scale-Fatigue (VAS-F)**, Lee et al (1991) was used to assess general fatigue, both physical and mental. The difference between pre- and post-task scores served as the primary index of task-induced fatigue.
**-** **NASA Task Load Index (NASA-TLX)**, Hart (2006) was used to assess workload after the task.

Participants rated their workload experience across six dimensions on a 100-point scale with 5-point increments. Raw scores were averaged across items due to increase their test-retest reliability (Bustamante and Spain, 2008), and differences between conditions were compared.

#### 2.5.2. Behavioural analysis

The accuracy and speed of participants’ responses to the double jump events were measured based on space bar key presses recorded during data collection. A failure to press the key within 3 seconds after a target event was classified as a miss. Any key press recorded between this 3-second window and the next target event was considered accidental and excluded from further analysis. To examine time-on-task effects, the number of missed target events and the median reaction time were calculated for each of the six consecutive time periods. These measures were compared across the two experimental sessions to evaluate the effect of flicker presence on task performance.

#### 2.5.3. SSVEP analysis

The continuous SSVEP response elicited by the 14Hz minimally intrusive flicker superimposed on the task was extracted using the Rhythmic Entrainment Source Separation (RESS; Cohen and Gulbinaite 2017) method. RESS computes a spatial filter that maximizes the signal-to-noise ratio (SNR) of the SSVEP response at a specific frequency. First, channel-to-channel covariance matrices were computed from narrow-band filtered data centered around the 14Hz stimulation frequency (using a Gaussian-shaped filter with a full-width half maximum (FWHM) of 1 Hz) as well as from neighboring frequencies. These neighboring frequencies (1 Hz away from the stimulation frequency, with a FWHM of 1 Hz) provided reference covariance matrices (R matrices). A generalized eigendecomposition was then applied between the stimulation frequency and the average of the neighboring frequencies’ covariance matrices. The eigenvector corresponding to the largest eigenvalue was selected as the main RESS component and back-projected onto the EEG time series. Since the RESS method optimizes the SNR at the stimulation frequency, caution is necessary when interpreting the filtered data, as overfitting is possible (Cohen and Gulbinaite 2017). To verify that the 14Hz RESS component was not merely a by-product of this optimization process, the 14Hz RESS component was also computed on EEG data recorded during the no-flicker condition.

#### 2.5.4. Alpha power analysis

Although historically regarded as a marker of an “idling brain,” recent research suggests a more nuanced role for alpha oscillations. Rhythmic activity around 10 Hz is now understood to facilitate effective communication between brain regions, playing a crucial role in inhibition and gating processes that suppress irrelevant activity. Rather than averaging across the entire alpha range (8–12 Hz), focusing on the individual alpha peak frequency (IAPF) provides a more sensitive measure of neural dynamics, such as processing speed (Klimesch et al 1996), and is linked to various cognitive functions (Haegens et al 2014; Hülsdünker et al 2016). As the IAPF can vary significantly across individuals (Grandy et al 2013), accounting for these individual differences is essential for more accurate analysis.

In this study, the IAPF was calculated from 150 seconds of resting-state data recorded before the experimental task, during which participants had their eyes closed. The IAPF was identified using the algorithm proposed by Corcoran et al (2018), with the recommended parameters (Fw = 11, k = 5). This individualized alpha frequency was then used to design filters for generalized eigendecomposition (GED), which enhances the contrast between signal components centered around the IAPF and broader, broadband activity. GED decomposes the data by comparing two covariance matrices: one representing the signal of interest (centered on the IAPF with a FWHM of 0.5 Hz) and the other representing broadband noise (within the 1 to 40 Hz bandpass-filtered signal) Parra and Sajda (2003). The eigenvector corresponding to the largest eigenvalue was selected as the primary component, effectively acting as a combined spectral and spatial filter (Cheveigné and Arzounian 2015). This primary component filter was applied to the continuous time series EEG data to retrieve alpha power fluctuations in the time domain (Cohen, 2022). The continuous filtered data was then segmented into epochs of 1, 3, 5, and 10 seconds preceding hits and misses trials. Mean alpha peak activity for each time window was extracted and compared across events.

#### 2.5.5. Theta power analysis

Frontal theta activity has been shown to gradually increase during prolonged tasks (Laukka et al 1995) and is positively associated with task demands (Onton et al, 2005). Its consistent relationship with time-on-task effects, regardless of modality or task, suggests it reflects the rising cost of mental fatigue (Strijkstra et al, 2003) and executive control (Wascher et al, 2014; Arnau et al, 2017), leading to its recognition as a marker of mental effort. In this study, midfrontal theta was isolated using spatial filters designed to maximize theta-band power. This approach is similar to the GED-based method used to extract alpha activity described in the previous section. Building on the method of Arnau et al (2017), we constructed the spatial filter by optimizing channel weights to maximize the distinction between theta-band (4–8 Hz) and broadband activity. A temporal filter centered around 6 Hz with a full-width half maximum (FWHM) of 2 Hz was applied, while broadband activity was filtered within the 1 to 40 Hz range. As with the alpha analysis, the filtered continuous data was segmented into epochs of 1, 3, 5, and 10 seconds preceding hits and misses trials. Mean theta activity for each time window was extracted and compared across events.

### 2.6. Statistical analyses

The primary objective of this study was to evaluate the effectiveness of a low-intensity flicker stimulus in eliciting SSVEP responses and to determine whether the amplitude of these responses could predict lapses of attention. To do so, we compared SSVEP amplitudes recorded during the periods preceding correct target detections (hits) and missed target detections (misses). In the first analysis, various time intervals preceding these events (1, 3, 5, and 10 seconds) were explored to identify the optimal temporal scale for detecting attentional lapses. A repeated measures Analysis of Variance (ANOVA) was conducted, with time window (1, 3, 5, and 10 seconds), experimental condition (flicker, no flicker), and trial outcome (hits, misses) as independent factors, to examine their effects on SSVEP amplitude. Similar analyses were conducted on theta and alpha power. Holm corrections for multiple comparisons were applied to all post-hoc paired-sample t-tests investigating main effects and interactions between the factors in the repeated measures ANOVAs. In cases where Mauchly’s test indicated a violation of the sphericity assumption, a Greenhouse-Geisser correction was applied.

The second aim of this study was to assess the impact of the flicker probe on subjective experience and task performance. Subjective measures of eye strain, fatigue, sleepiness, and mental workload, along with hit rate and median reaction time to target stimuli, were compared between the flicker and no-flicker conditions. For this analysis, two-tailed paired-samples t-tests were performed on task performance measures and the raw scores of the NASA-TLX questionnaire, as well as the pre-post task difference scores of other scales. A Shapiro-Wilk test of normality was conducted for each measure. If the assumption of normality was violated, Wilcoxon signed-rank tests were used in place of paired-samples t-tests.

## 3. Results

### 3.1. User experience

The differences in terms of user experience between the flicker and no flicker sessions were investigated across four dimensions: sleepiness, fatigue, eye strain, and perceived task load. The comparison of the pre-post difference scores on the Karolinska Sleepiness Scale did not reveal significant difference in sleepiness between the flicker and no flicker conditions [t(15) = .513, p > .05, d = .128]. The subjective assessment of fatigue induced by the performance of the task did not significantly vary across conditions [W = 61.00, z = .362, p > .05, corr = - .103]. Similarly, the DES questionnaires administered during both sessions did not reveal a significant difference related to the presence of a flicker in terms of eye strain [W = 34.5, z = .714, p > .05, corr = .255]. The perceived task load assessed through the NASA-TLX questionnaire was similar across conditions [t(15) = .068, p > .05, d = .017]. A summary of the data can be visualized in Figure 2A-D.

**Figure 2:**
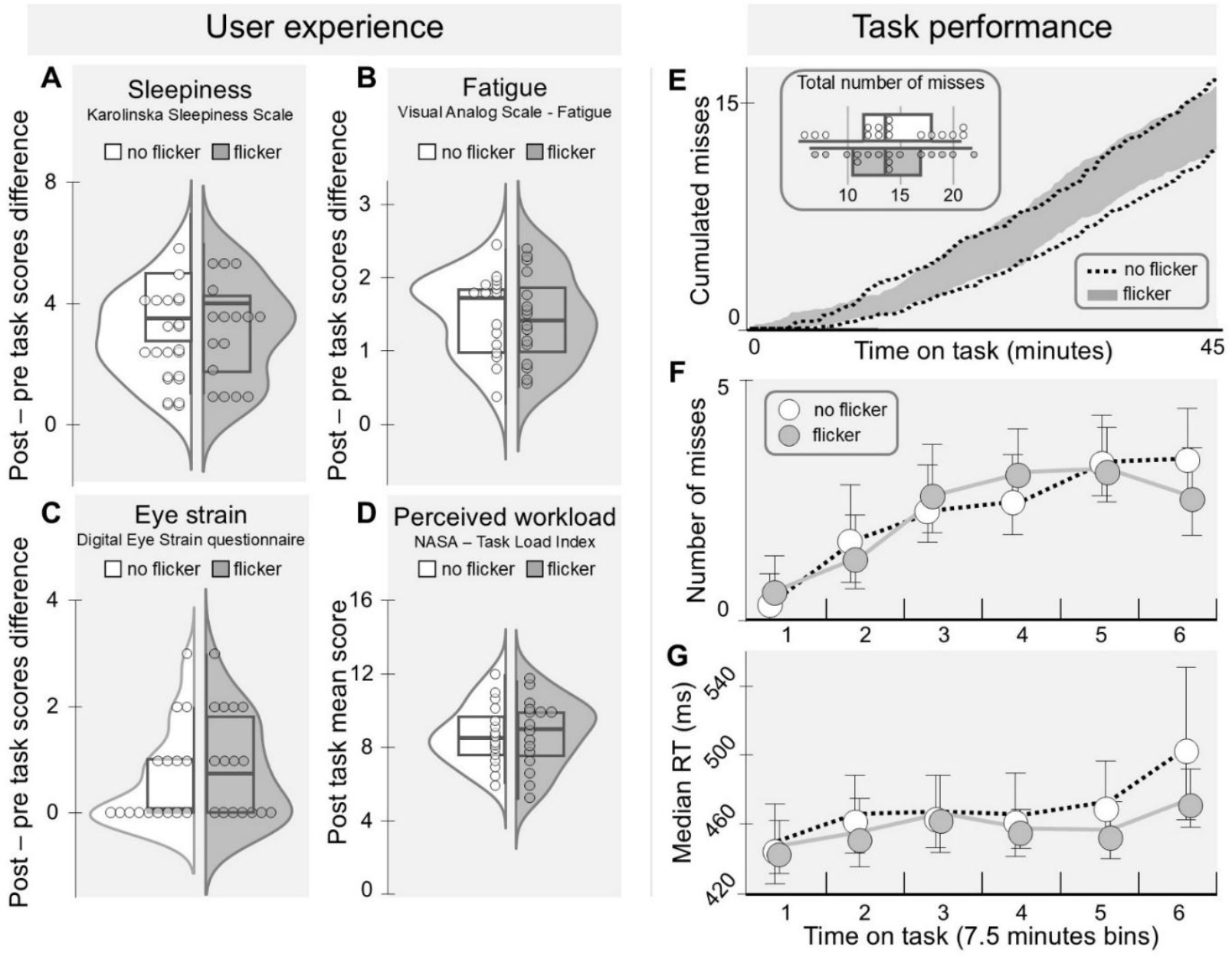
Comparison of user experience and task performance measures between the experimental conditions (with or without the 14 Hz transparent flicker) recorded in separate sessions. Differences between pre- and post-task measures of sleepiness (A), fatigue (B), and eye strain (C) were compared between sessions (no flicker in white, with flicker in gray) to evaluate the impact of the presence of the transparent flicker on participants’ subjective experiences. Perceived workload (D) was measured at the end of each session. Task performance measures, including the cumulative number of missed events over time-on-task and total number of misses per session (E), as well as the median number of missed events (F) and median reaction times (G) per 7.5-minute period, were compared across conditions to assess time-on-task effects.

### 3.2. Behavioural performance

Task performance was assessed in terms of reaction time and number of missed targets. No difference was found between the flicker and control conditions in terms of median reaction time [W = 47, z = 1.086, p = .298, corr. = 1.294] and total number of missed targets [t(1,15) = .09, p > .05, d = .023]. These results suggest that the presence of the flicker did not affect task performance. In order to investigate whether measures reflecting the time on task effect were affected by the presence of a flicker, the continuous data was split into six consecutive windows of 7.5 minutes, from which outcome measures were extracted. The behavioural measures (median reaction time and number of errors per period) were analyzed through 2-by-6 repeated-measures ANOVAs with task on time (first to sixth time period) and condition (flicker, no flicker) as main factors. There was no effect of the condition (presence or absence of the transparent flicker) on the number of missed events [F(1,15) = 0.051, p > .05,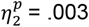]. The time on task however had a main effect on the number of missed events [F(5,75) = 15.699, p < .001,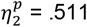]. The interaction between the two factors did not yield a main effect on the number of errors commited [F(5,75) = 0.786, p > .05,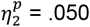]. Post-hoc test showed that a significantly lower number of events were missed during the first two periods (from 0 to 15 minutes into the task) than over the rest of the task (see Figure 2E-F and Figure 5). There was no significant difference in terms of number of missed events between the following periods.

Similarly, reaction time was not found to be mainly affected by the presence or absence of the transparent flicker [F(1,15) = 0.933, p > .05,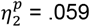]. A main effect of the time on task was also observed on reaction time [F(1.723,25.838) = 4.896, p < .05,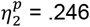]. The interaction of the two factors did not alter the reaction time significantly [F(2.237,33.559) = 0.726, p > .05,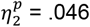]. Post-hoc paired-sample t-tests revealed that reaction time significantly increased during the last period (37.5 to 45 minutes) when compared to the first and second time periods (from 0 to 15 minutes into the task) [t(1,15) = 4.764, p < .001, d = .531 and t(1,15) = 3.363, p < .05, d = .375, respectively]. No significant difference in reaction time was observed between the first two periods, nor between the following periods when compared amongst themselves (see Figure 2G).

### 3.3. EEG analysis

This study aimed to evaluate the validity of a continuous yet minimally intrusive SSVEP frequency-tagging approach for measuring lapses of attention during a sustained visual attention task. SSVEP measures were compared with other commonly used markers of attention and vigilance, including pre-stimulus theta and alpha band power. As a first validation step, we examined the effects of flicker presence and time-on-task on these EEG measures to assess both their reliability and potential interference with the proposed approach. This analysis was performed on the median signal features computed across six consecutive time periods of 7.5 minutes, comparing the two experimental sessions where participants performed the clock task with and without the 14 Hz flicker. Next, to determine which EEG features could differentiate between successful target detections (hit trials) and missed detections (miss trials), EEG features preceding these events were extracted from the continuous signals recorded during the flicker condition and compared. Additionally, to identify the optimal time window for distinguishing hit and miss trials using band-specific features, spectral power measures were computed for varying temporal windows (1, 3, 5, 10, and 30 seconds) preceding each event.

#### 3.3.1. Steady-States Visually Evoked Potentials (SSVEP)

As expected, the presence of the transparent flicker had a main effect on the strength of the SSVEP signal extracted [F(1,15) = 16.454, p < .001,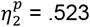]. The time on task also had a main effect on SSVEP response [F(5, 75) = 2.809, p < .05,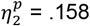]. The interaction between flicker presence and time on task yielded a main effect on SSVEP response [F(5, 75) = 4.661, p < .001,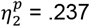]. Post-hoc tests revealed that the strength of the SSVEP signal was higher when the flicker was present than absent [t(1,15) = 3.390, p < .001, d = .736]. Moreover, the amplitude of the SSVEP response elicited by the presence of the transparent flicker was significantly higher at the end of the task than at its start (see Figure 3A).

**Figure 3:**
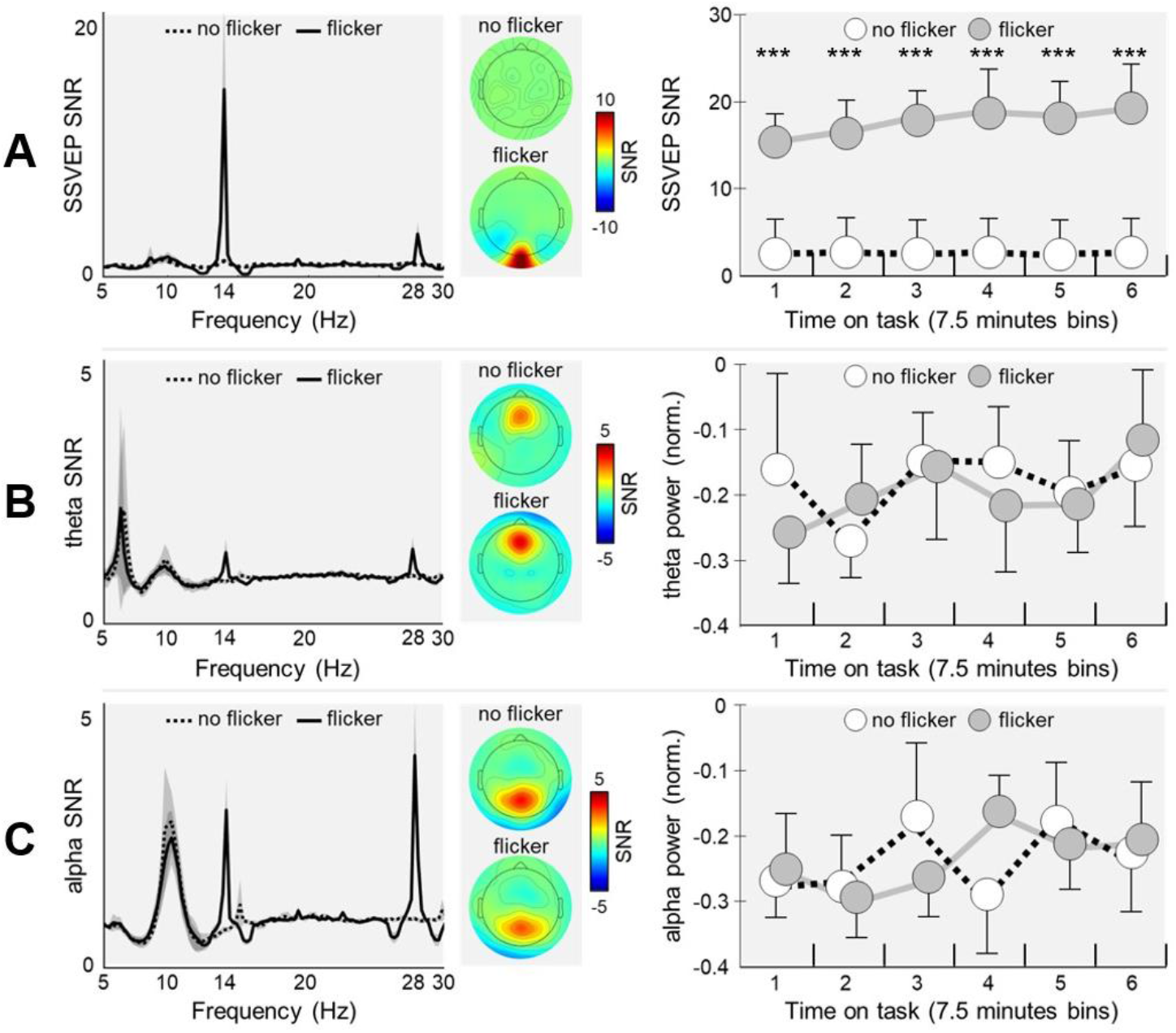
Grand-average (N=16) power spectral density, topography, and time-on-task modulations of three EEG features of interest: SSVEP response (A), theta power (B), and alpha peak power (C) across two experimental sessions (with and without a 14 Hz transparent flicker). Left panels: Power spectral density of the spatially filtered signals recorded over the entire 45-minute task. Solid lines represent the flicker condition; dotted lines represent the no-flicker condition. Center panels: Mean topographical distribution of the spatial filters associated with each signal of interest, shown separately for both flicker and no-flicker conditions. Right panels: Fluctuations of each signal across the 45-minute task, binned into six consecutive 7.5-minute segments to evaluate the impact of flicker presence and time-on-task. Pairwise comparisons (t-tests) between flicker conditions that reached significance are denoted by asterisks (p <.001***).

The detection of the event (hits or miss) had a main effect on SSVEP signal strength [F(1, 15) = 148.35, p < .001,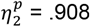]. The temporal window length over which the mean signal was extracted had a main effect on SSVEP power [F(1.764, 26.455) = 29.732, p < .001,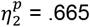]. The interaction between window length and detection factors affected the amplitude of the SSVEP [F(1.437, 21.559) = 53.918, p < .001,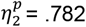]. Post-hoc tests revealed that, across time windows of different lengths, the SSVEP signal strength was significantly higher when participants detected the target events than when they missed them [t(15) = 12.180, p < .001, d = 4.069]. More precisely, these analyses revealed that a one second time window led to the highest contrast between SSVEP signals preceding hits and misses [t(15)= 15.507, p < .001, d = 6.475] (see Figure 4A), although the SSVEP response over three [t(15) = 12.300, p < .001, d = 5.136], five [t(15) = 10.656, p < .001, d = 4.450], and ten [t(15) = 10.762, p < .001, d = 3.241] seconds were also significantly different between hits and misses trials. This higher contrast for the one second window is mainly explained by a significant reduction in SSVEP signal amplitude during the last second preceding the missed target events than with using any of the longer time windows whereas the amplitude of the SSVEP response preceding the successfully detected target events did not vary across time windows (see Figure 4D). The extraction of mean SSVEP signal over a 30 second window preceding the events did not lead to a significant difference between hits and misses trials [t(15) = 2.495, p > .05, d = 1.042].

**Figure 4:**
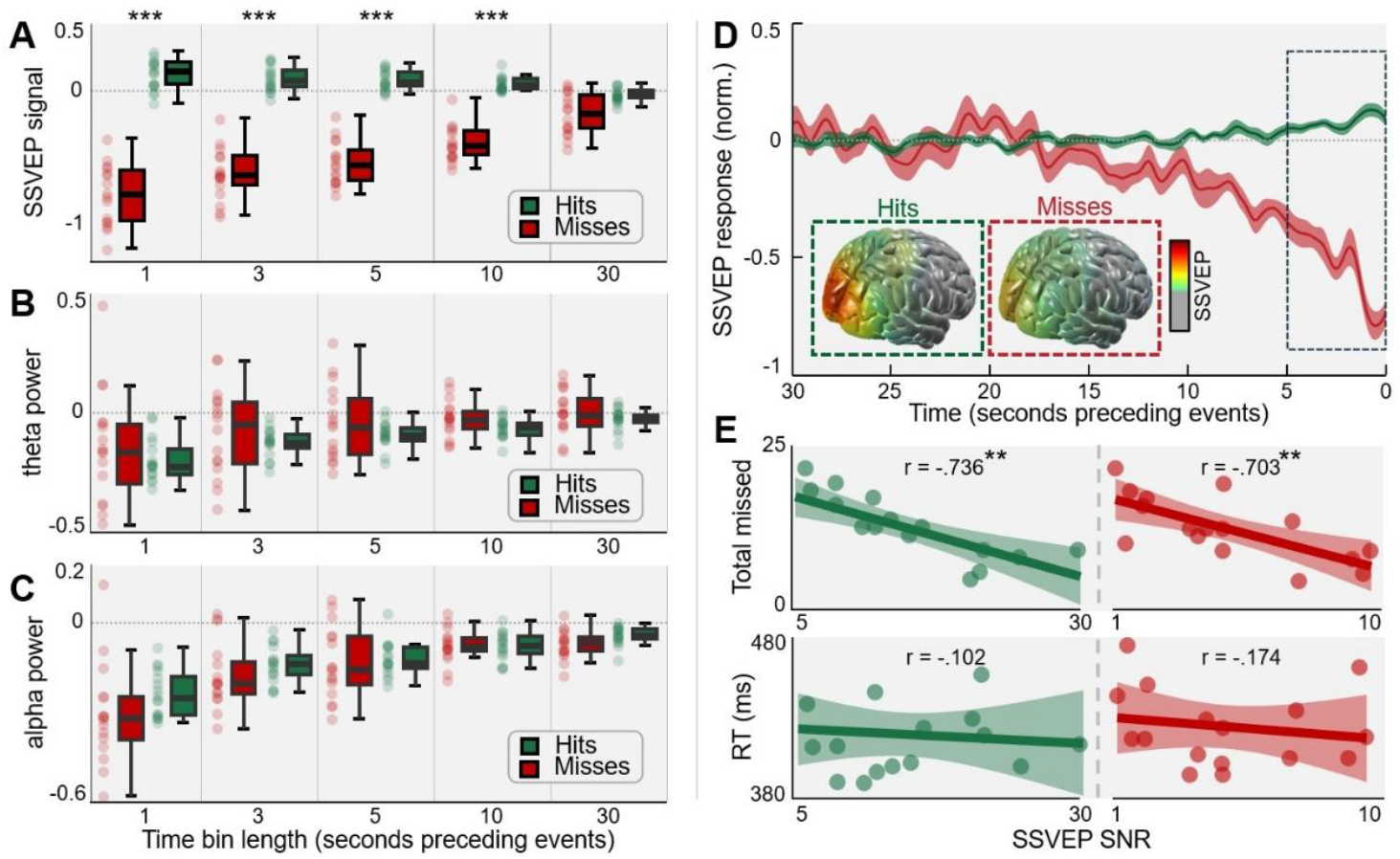
Distributions of mean SSVEP responses (A), theta power (B), and alpha power (C) in the 1, 3, 5, 10, and 30 seconds preceding hit (green) and missed (red) events. D. Grand average (N=16) SSVEP response waveforms preceding hit and missed events. The dotted insets show topographical maps depicting correlations between (1) the forward model of the main SSVEP component (extracted from the final five seconds preceding hit and missed events) and (2) a leadfield matrix based on standard MRI and BEM models (Cohen & Gulbinaite, 2017). We used the median correlation across all voxels as a threshold: subthreshold correlation values appear in gray, whereas values exceeding this threshold are depicted by a color map ranging from green to red (maximal correlation). (E) Correlations between the mean SSVEP response recorded one second before hit (green) and missed (red) events with the total number of missed trials and the median reaction time (RT) across participants. Pairwise comparisons (t-tests) between hit and missed events across temporal windows and correlations that reached significance are marked by asterisks (p < .01**, p < .001***).

**Figure 5:**
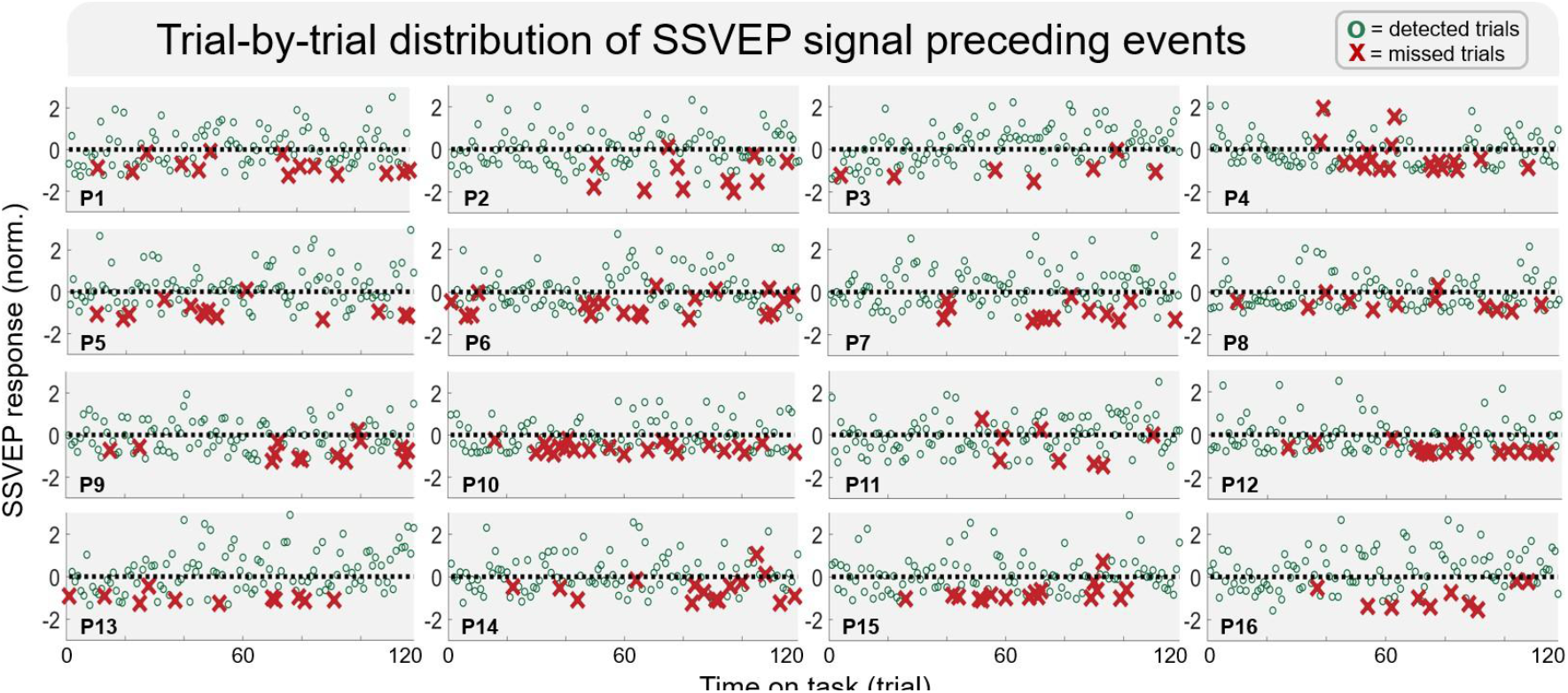
Subject-specific trial-by-trial fluctuations in SSVEP responses recorded one second before target events during the 45-minute clock task, with a transparent flicker superimposed. Green circles represent SSVEP responses recorded prior to detected trials, while red crosses represent responses preceding missed trials. The SSVEP responses were z-score normalized based on the entire recording. The dotted line represents the mean SSVEP response across the 45-minute task.

#### 3.3.2. Theta power

The presence of the flicker did not have a main effect on overall theta signal strength [F(1,15) = 0.288, p > .05,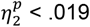]. There was no main effect of time on task on theta power [F(3.014, 45.207) = 1.372, p > .05,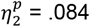]. The interaction between flicker presence and time on task did not yield a main effect on theta power [F(5, 75) = 1.284, p > .05,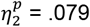]. These results indicate that neither the presence of the transparent flicker nor the time on task did have an effect on overall theta power (see Figure 3B).

The detection of the target event (hits or miss) did not have a main effect on theta power [F(1, 15) = 1.846, p > .05,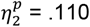]. The length of the temporal window over which the mean signal was extracted did not have a main effect on theta power [F(2.570, 38.554) = 12.023, p < .001,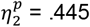]. The interaction between temporal window length and target event detection factors did not affect the theta signal recorded [F(2.458, 36.876) = 0.079, p > .05,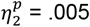] (see Figure 4B).

#### 3.3.3. Alpha peak power

The presence of the flicker did not have a main effect on overall alpha signal strength [F(1,15) = 0.012, p > .05,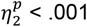]. There was no main effect of time on task on alpha peak signal [F(5, 75) = 1.666, p > .05,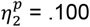]. Moreover, the interaction between flicker presence and time on task did not affect alpha peak power [F(5, 75) = 2.204, p > .05,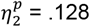]. These results indicate that neither the presence of the flicker nor the time on task influenced the alpha peak signal strength (see Figures 3C).

The detection of the target event (hits or miss) did not have a main effect on alpha peak signal strength [F(1, 15) = 1.392, p > .05,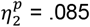]. By contrast, the length of the temporal window over which the mean signal was extracted did not have a main effect on signal strength [F(2.589, 38.838) = 34.705, p < .001,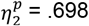]. The interaction between temporal window length and target event detection factors did not affect the alpha peak signal recorded [F(2.437, 36.550) = 1.217, p > .05,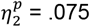] (see Figure 4C).

#### 3.3.4. Correlation between task performance and EEG features

The SSVEP signal amplitude preceding both the detected and the overlooked target events were negatively correlated with the total number of missed target events [detected: r(15) = -.736, p < .01, z = - .942; undetected: r(15) = -.703, p < .01, z = -.873]. There was however no significant relationship between the median reaction time and neither the SSVEP amplitude preceding detected [r(15) = -.102, p > .05] and undetected target events [r(15) = -.174, p > .05] (see Figure 4E). Moreover, the SSVEP signal amplitude preceding detected and overlooked target events was strongly correlated [r(15) = .870, p < .001, z = 1.335]. For the other frequency band features, the median reaction time was not significantly correlated with peak alpha signal preceding hits [r(15) = .206, p > .05] and missed trials [r(15) = -.171, p > .05]. There was no significant relationship between the total number of errors and peak alpha signal recorded before the occurrence of hits [r(15) = .118, p > .05] but with missed trials [r(15) = .537, p < .05, z = .600]. The measures of task performance were not significantly correlated with theta activity recorded before the detection or oversight of target events.

## 4. Discussion

This study investigated whether a minimally intrusive SSVEP frequency-tagging approach could effectively monitor vigilance fluctuations and predict lapses of attention during a prolonged sustained attention task. A 14 Hz low-luminance, low-contrast flicker was superimposed onto the Mackworth Clock Task, a well-established paradigm for simulating real-world monitoring scenarios. This design aimed to capture reliable neural markers of attentional lapses while minimizing sensory disruption. A control condition, in which participants performed the task without the flicker, confirmed that SSVEP measures were specifically modulated by the flicker without affecting user experience or task performance.

### 4.1. Behavioral Performance and Subjective Experience

The primary aim of our analyses was to verify whether the experimental paradigm effectively induced attentional lapses. The behavioral data clearly demonstrated a time-on-task effect: as the task progressed, participants’ performance deteriorated, evidenced by an increase in missed target events and slower reaction times. These findings align with established research on vigilance lapses (Mackworth, 1948; Martel et al 2014; Durantin et al 2015; Braboszcz and Delorme, 2011; Manly et al 1999; Robertson et al 1997; Drummond et al 2005), confirming that sustained attention tasks lead to performance declines over time. Importantly, the presence of the low-intensity flicker did not affect task performance. Reaction times and accuracy were comparable between the control condition (without flicker) and the experimental condition (with flicker), indicating that the flicker did not interfere with participants’ ability to perform the task.

Subjective reports further supported these findings. The contrast between pre- and post-task self-reports of sleepiness, eye strain, and fatigue did not differ between the flicker and non-flicker conditions, suggesting that the flicker did not negatively impact participants’ experience. Moreover, participants’ perception of task difficulty remained consistent across both conditions, reinforcing the minimal effect of the flicker on task performance and user comfort. These results are particularly relevant given the well-documented issues with traditional flicker stimuli, which are often associated with visual discomfort and distraction (Zhu et al 2010; Makri et al 2015; Cao et al 2014; Ortner et al 2011; Patterson Gentile and Aguirre 2020).

### 4.2. Neural Response and SSVEP Modulation

While subjective assessments confirmed that the flicker was visually comfortable and did not impair task performance, it was critical to assess whether it could elicit measurable neural responses in the visual cortex and track fluctuations in vigilance, potentially predicting attentional lapses. Given the reduced saliency of the stimuli, it was initially anticipated that detecting neural variations might be challenging. However, signal-to-noise ratio (SNR) analyses revealed that even low-contrast flickers successfully triggered cortical responses, as demonstrated by measurable SNR values.

Moreover, the SSVEP signal at the target frequency was significantly modulated by the presence of the subtle flicker, increasing monotonically with time-on-task. Post-hoc analyses confirmed that SSVEP amplitude was significantly higher at the end of the task than at its onset, reinforcing this trend. These findings may seem counterintuitive, particularly in light of previous research showing a reduction in SSVEP SNR with prolonged task engagement and presumed mental fatigue (Lamti et al 2014; Xie et al 2016). However, it is important to note that in these studies, participants were not engaged in a vigilance task but were focused solely on the flicker stimuli, which likely induced visual fatigue and eye strain, rather than mental fatigue.

An alternative explanation for our results is that prolonged time-on-task induces mental fatigue, which paradoxically triggers stronger neural responses as a compensatory mechanism to offset reduced cognitive efficiency. This heightened neural activity could explain the increased SSVEP SNR observed in our study. Indeed, several studies suggest the existence of compensatory neural mechanisms that counteract cognitive fatigue. For example, Wang et al (2016) found that during the initial phase of a task (lasting up to approximately 80 minutes, termed the “compensation phase”), the brain adapts to cognitive fatigue by increasing neural activity. Beyond this phase, in the “decompensation phase,” the brain’s ability to compensate diminishes as fatigue intensifies. This compensation model is relevant to our study, as the task duration of approximately 40 minutes aligns with the compensation phase, where compensatory mechanisms are likely to be active. The lack of changes in traditional markers such as alpha and theta rhythms which are generally known to index time-on-task effects, as reported in several studies (Groot et al 2021; Molina et al 2019; Dinges and Powell 1985; Drummond et al 2005; Esposito et al 2022) may further suggest that the task duration was insufficient to trigger the decompensation phase. This, in turn, supports the hypothesis that SSVEP may serve as a more sensitive indicator of attentional fluctuations compared to other neural markers.

Another possible explanation is that, during prolonged tasks, the brain may adapt to the flicker stimulus, amplifying the SSVEP response to maintain vigilance. This adaptation could involve greater engagement of the occipital cortex, as participants increase their effort to remain attentive, reflecting an adaptive neural strategy. Future studies should explore longer task durations as in (Boksem et al 2005; Wascher et al 2014), potentially using the Mackworth Clock Test, to further investigate these hypotheses. This would clarify the impact of extended time-on-task on SSVEP SNR under conditions of prolonged fatigue and provide deeper insights into the role of compensatory mechanisms in sustaining attention.

### 4.3. SSVEP Response as a Predictor of Attentional Lapses

An additional objective of this study was to determine whether steady-state visual evoked potentials (SSVEP) could act as sensitive markers for predicting lapses in attention. Our findings reveal that the SSVEP signal-to-noise ratio (SNR) was significantly lower in the moments preceding missed target events compared to those correctly detected. This result suggests that SSVEP amplitude may serve as a reliable indicator of vigilance, with the potential to distinguish between episodes of inattention and effective attentional engagement as early as 15 seconds before a response. Notably, shorter time windows (e.g., one second) prior to target events yielded the most robust discrimination between hits and misses, highlighting the temporal precision of SSVEP as a continuous measure of attention. Further correlational analyses reinforced the predictive value of SSVEP, as these measures were linked to both correct detections and misses. Together, these results indicate that SSVEP is not only sensitive to attentional lapses but also functions as a reliable predictor of performance outcomes.

A potential confound considered was the possibility that these SSVEP changes could reflect shifts in endogenous brain rhythms, particularly the alpha rhythm (around 14 Hz), which is close to the flicker frequency. However, no such modulations were observed in a control condition without flicker stimulation, ruling out the influence of endogenous alpha activity and confirming that the observed SSVEP variations were specific to the flicker stimulus. Moreover, both alpha and theta band activity did not show significant differences between hit and miss trials during event detection as previously shown by Shou et al (2015); Martel et al (2014); O’Connell et al (2009). Nevertheless, further analyses revealed a positive correlation between alpha power and missed trials. This finding aligns with existing literature, where elevated alpha activity is often associated with attentional disengagement and performance lapses (Groot et al 2021; Molina et al 2019; Dinges and Powell 1985; Drummond et al 2005; Esposito et al 2022).

However, alpha power was not correlated with hit trials, suggesting that SSVEP provides a more specific predictor of attentional engagement. Overall, these results underscore the utility of SSVEP as a distinct and reliable indicator of attention that is not confounded by endogenous rhythms such as alpha, which remained unaffected by flicker in control conditions. Taken together, the findings highlight the potential of SSVEP for long-term vigilance monitoring, offering advantages over traditional spectral markers like theta and alpha activity. In particular, the lack of temporal stability and specificity of these traditional markers, especially in real-world applications, reinforces the superiority of SSVEP as a tool for continuous vigilance assessment. These findings provide a strong basis for the application of SSVEP in monitoring sustained attention across various contexts.

### 4.4. Limitations and Future Directions

While our findings demonstrate the efficacy of SSVEP for detecting attentional lapses, several limitations should be addressed in future research. First, our sample size of 16 participants, though sufficient for exploratory purposes, may limit the generalizability of the results. Larger and more diverse samples are needed to ensure the robustness of these findings. Additionally, this study used a single flicker frequency (14 Hz) to avoid interference with endogenous alpha rhythms. Future work could explore the efficacy of other flicker frequencies and modulation depths to optimize both user comfort and signal reliability.

Moreover, while our study focused primarily on neural markers, integrating additional behavioral data such as eye-tracking measures could provide a more comprehensive understanding of attentional dynamics. Combining SSVEP with other behavioral indices may offer deeper insights into how neural activity corresponds to task performance.

### 4.5. Conclusion

In conclusion, this study demonstrates the effectiveness of a minimally intrusive SSVEP frequency-tagging approach for real-time monitoring of vigilance decrements. SSVEP responses, triggered by a low-luminance, low-contrast 14 Hz flicker, accurately predicted lapses in attention and outperformed traditional alpha-band EEG markers. Crucially, the flicker stimulus did not disrupt task performance or user comfort, making it highly suitable for high-stakes industries like aviation and healthcare. These findings pave the way for developing human-computer interfaces (HCIs) capable of continuously monitoring cognitive states in environments where attentional lapses could have critical consequences. By detecting vigilance lapses in real time, such HCIs could trigger adaptive interventions, such as task reminders or automated assistance, to maintain optimal performance.

## Acknowledgment

This work was funded by the European Community and DGAC (CORAC/Toucans project), the AXA Research fund, and ANITI (Chair for Neuroadaptive Technology).

## Ethical statement

This study was approved by the Ethics Committee of the University of Toulouse (CER approval number 2023-749) and conducted in accordance with the Declaration of Helsinki. Written informed consent was obtained from all participants before the experiment.

## Data and Code Availability

The datasets analyzed in the current study will be made publicly available in an online repository. The source code for the analysis will also be available on GitHub.

## CRediT authorship contribution statement

**S. Ladouce:** Conceptualization, Data collection, Data analysis, Writing. **J. J. Torre-Tresols:** Data analysis, Writing. **K. Le Goff:** Writing. **F Dehais:** Conceptualization, Writing.

## Footnotes

1 A video showcasing the implementation of the transparent flicker stimulus can be accessed at the following address: https://www.youtube.com/watch?v=GELUF-WA8xk

## References

Adamian, N., & Andersen, S. K. (2024). Attentional Modulation in Early Visual Cortex: A Focused Reanalysis of Steady-state Visual Evoked Potential Studies. Journal of Cognitive Neuroscience, 36(1), 46–70. 10.1162/jocn_a_02070 Akerstedt, T., Gillberg, M., 1990. Subjective and objective sleepiness in the active individual. The International journal of neuroscience 52, 29–37. doi:10.3109/00207459008994241.

Andersen, S. K., Müller, M. M., & Hillyard, S. A. (2012). Tracking the allocation of attention in visual scenes with steady-state evoked potentials. In Cognitive neuroscience of attention, 2nd ed. (pp. 197–216). The Guilford Press.

Andrillon, T., Burns, A., Mackay, T., Windt, J., Tsuchiya, N., 2021. Predicting lapses of attention with sleep-like slow waves. Nature Communications 12, 1–12. URL: http://dx.doi.org/10.1038/s41467-021-23890-7, doi:10.1038/s41467-021-23890-7.

Arnau, S., Möckel, T., Rinkenauer, G., Wascher, E., 2017. The interconnection of mental fatigue and aging: An EEG study. International Journal of Psychophysiology 117, 17–25. doi:10.1016/j.ijpsycho.2017.04.003.

Arora, K., Gayet, S., Kenemans, J.L., der Stigchel, S.V., Chota, S., 2024. Rapid Invisible Frequency Tagging (RIFT) in a novel setup with EEG. bioRxiv URL: https://www.biorxiv.org/content/early/2024/02/04/2024.02.01.578462, doi:10.1101/2024.02.01.578462.

Boksem, M.A., Meijman, T.F., Lorist, M.M., 2005. Effects of mental fatigue on attention: An ERP study. Cognitive Brain Research 25, 107–116. doi:10.1016/j.cogbrainres.2005.04.011.

Boremanse, A., Norcia, A.M., Rossion, B., 2013. An objective signature for visual binding of face parts in the human brain. Journal of Vision 13, 6–6.

Borghini, G., Astolfi, L., Vecchiato, G., Mattia, D., Babiloni, F., 2014. Measuring neurophysiological signals in aircraft pilots and car drivers for the assessment of mental workload, fatigue and drowsiness. Neuroscience & Biobehavioral Reviews 44, 58–75.

Braboszcz, C., Delorme, A., 2011. Lost in thoughts: Neural markers of low alertness during mind wandering. NeuroImage 54, 3040–3047. doi:10.1016/j.neuroimage.2010.10.008.

Brosowsky, N.P., DeGutis, J., Esterman, M., Smilek, D., Seli, P., 2023. Mind Wandering, Motivation, and Task Performance Over Time: Evidence That Motivation Insulates People From the Negative Effects of Mind Wandering. Psychology of Consciousness: Theory Research, and Practice doi:10.1037/cns0000263.

Bustamante, E.A., Spain, R.D., 2008. Measurement Invariance of the Nasa TLX. Proceedings of the Human Factors and Ergonomics Society Annual Meeting 52, 1522–1526. URL: https://doi.org/10.1177/154193120805201946, doi:10.1177/154193120805201946.

Cabrera-Castillos, K., Ladouce, S., Darmet, L., Dehais, F., 2023. Burst c-VEP Based BCI: Optimizing stimulus design for enhanced classification with minimal calibration data and improved user experience. NeuroImage 284, 120446. doi:10.1016/j.neuroimage.2023.120446

Cao, T., Wan, F., Wong, C.M., da Cruz, J.N., Hu, Y., 2014. Objective evaluation of fatigue by EEG spectral analysis in steady-state visual evoked potential-based brain-computer interfaces. BioMedical Engineering Online 13, pp. 1–13. doi:10.1186/1475-925X-13-28.

Chen, X., Wang, Y., Zhang, S., Xu, S., Gao, X., 2019. Effects of stimulation frequency and stimulation waveform on steady state visual evoked potentials using computer monitor. Journal of Neural Engineering 16. doi:10.1088/1741-2552/ab2b7d.

Cheveigné, A.D., Arzounian, D., 2015. Scanning for oscillations. Journal of Neural Engineering 12, 066020. doi:10.1088/1741-2560/12/6/066020.

Chun, M.M., Golomb, J.D., Turk-Browne, N.B., 2011. A Taxonomy of external and internal attention. Annual Review of Psychology 62, 73–101. doi:10.1146/annurev.psych.093008.100427.

Clayton, M.S., Yeung, N., Cohen Kadosh, R., 2015. The roles of cortical oscillations in sustained attention. doi:10.1016/j.tics.2015.02.004.

Cohen, M., Gulbinaite, R., 2017. Rhythmic entrainment source separation: Optimizing analyses of neural responses to rhythmic sensory stimulation. NeuroImage 147. doi:10.1016/j.neuroimage.2016.11.036.

Cohen, M.X., 2022. A tutorial on generalized eigendecomposition for denoising, contrast enhancement, and dimension reduction in multi-channel electrophysiology. NeuroImage 247, 118809. URL: https://doi.org/10.1016/j.neuroimage.2021.118809, doi:10.1016/j.neuroimage.2021.118809.

Corcoran, A.W., Alday, P.M., Schlesewsky, M., Bornkessel-Schlesewsky, I., 2018. Toward a reliable, automated method of individual alpha frequency (IAF) quantification. Psychophysiology 55, e13064. doi:10.1111/psyp.13064.

Czisch, M., Wehrle, R., Harsay, H.A., Wetter, T.C., Holsboer, F., Sämann, P.G., Drummond, S.P., 2012. On the need of objective vigilance monitoring: Effects of sleep loss on target detection and task-negative activity using combined EEG/fMRI. Frontiers in Neurology APR, 1–12. doi:10.3389/fneur.2012.00067.

Dehais, F., Lafont, A., Roy, R., Fairclough, S., 2020. A neuroergonomics approach to mental workload, engagement and human performance. Frontiers in neuroscience 14, 268.

Delorme, A., Palmer, J., Onton, J., Oostenveld, R., & Makeig, S. (2012). Independent EEG sources are dipolar. PLoS One, 7, e30135. x10.1371/journal.pone.0030135.

Delorme, A., Makeig, S., 2023. This is no “ICA bug”: response to the article, “ICA’s bug: how ghost ICs emerge from effective rank deficiency caused by EEG electrode interpolation and incorrect re-referencing”. Frontiers in Neuroimaging 2. doi:10.3389/fnimg.2023.1331404.

Ding, J., Sperling, G., & Srinivasan, R. (2006). Attentional modulation of SSVEP power depends on the network tagged by the flicker frequency. Cerebral Cortex (New York, N.Y. : 1991), 16(7), 1016–1029. 10.1093/cercor/bhj044 Dinges D.F., Powell, J.W., 1985. Microcomputer analyses of performance on a portable, simple visual RT task during sustained operations. Behavior Research Methods, Instruments, Computers doi:10.3758/BF03200977.

Drijvers, L., Jensen, O., Spaak, E., 2021. Rapid invisible frequency tagging reveals nonlinear integration of auditory and visual information. Human Brain Mapping 42, 1138–1152.

Drummond, S.P., Bischoff-Grethe, A., Dinges, D.F., Ayalon, L., Mednick, S.C., Meloy, M., 2005. The neural basis of the psychomotor vigilance task. Sleep 28, 1059–1068.

Durantin, G., Dehais, F., Delorme, A., 2015. Characterization of mind wandering using fnirs. Frontiers in systems neuroscience 9, 45.

Duszyk, A., Bierzyńska, M., Radzikowska, Z., Milanowski, P., Kus, R., Suffczyński, P., Michalska, M., Labeçki, M., Zwoliński, P., Durka, P., 2014. Towards an optimization of stimulus parameters for brain-computer interfaces based on steady state visual evoked potentials. PLoS ONE 9, 1–11. doi:10.1371/journal.pone.0112099.

Eichele, H., Juvodden, H.T., Ullsperger, M., Eichele, T., 2010. Mal-adaptation of event-related EEG responses preceding performance errors. Frontiers in Human Neuroscience doi:10.3389/fnhum.2010.00065.

Esposito, A., Braccili, E., Sgro, F., Chiarantano, E., D’Ippolito, M., Pisotta, I., Bigioni, A., Guerrieri, A., Mattia, D., Cincotti, F., 2022. Attention, Boredom and Mind Wandering during a Vigilance Task: EEG and Ocular Markers. 2022 IEEE International Workshop on Metrology for Extended Reality, Artificial Intelligence and Neural Engineering, MetroXRAINE 2022 - Proceedings, 477–482. doi:10.1109/MetroXRAINE54828.2022.9967678.

Esterman, M., Grosso, M., Liu, G., Mitko, A., Morris, R., DeGutis, J., 2016. Anticipation of monetary reward can attenuate the vigilance decrement. PLoS ONE doi:10.1371/journal.pone.0159741.

Gibbings, A., Ray, L.B., Berberian, N., Nguyen, T., Shahidi Zandi, A., Owen, A.M., Comeau, F.J., Fogel, S.M., 2021. EEG and behavioural correlates of mild sleep deprivation and vigilance. Clinical Neurophysiology 132, 45–55. doi: https://doi.org/10.1016/j.clinph.2020.10.010, doi:10.1016/j.clinph.2020.10.010.

Grandy, T.H., Werkle-Bergner, M., Chicherio, C., Lövdén, M., Schmiedek, F., Lindenberger, U., 2013. Individual alpha peak frequency is related to latent factors of general cognitive abilities. NeuroImage. doi:10.1016/j.neuroimage.2013.04.059.

Groot, J.M., Boayue, N.M., Csifcsák, G., Boekel, W., Huster, R., Forstmann, B.U., Mittner, M., 2021. Probing the neural signature of mind wandering with simultaneous fMRI-EEG and pupillometry. NeuroImage. doi:10.1016/j.neuroimage.2020.117412.

Gulbinaite, R., Van Viegen, T., Wieling, M., Cohen, M. X., & Vanrullen, R. (2017). Individual alpha peak frequency predicts 10 hz flicker effects on selective attention. Journal of Neuroscience, 37(42), 10173–10184. doi: 10.1523/JNEUROSCI.1163-17.2017

Gulbinaite, R., Roozendaal, D.H., VanRullen, R., 2019. Attention differentially modulates the amplitude of resonance frequencies in the visual cortex. NeuroImage 203, 116146.

Haegens, S., Cousijn, H., Wallis, G., Harrison, P.J., Nobre, A.C., 2014. Inter- and intra-individual variability in alpha peak frequency. NeuroImage doi:10.1016/j.neuroimage.2014.01.049.

Hancock, P.A., 2017. On the nature of vigilance. Human factors 59, 35–43.

Hart, S.G., 2006. NASA-task load index (NASA-TLX); 20 years later. Proceedings of the Human Factors and Ergonomics Society, 904–908doi:10.1177/154193120605000909.

Hillyard, S. A., Hinrichs, H., Tempelmann, C., Morgan, S. T., Hansen, J. C., Scheich, H., & Heinze, H. J. (1997). Combining steady-state visual evoked potentials and f MRI to localize brain activity during selective attention. Human Brain Mapping, 5(4), 287–292.

Hudson, A.N., Van Dongen, H.P., Honn, K.A., 2020. Sleep deprivation, vigilant attention, and brain function: a review. doi:10.1038/s41386-019-0432-6.

Hülsdünker, T., Mierau, A., Strüder, H.K., 2016. Higher balance task demands are associated with an increase in individual alpha peak frequency. Frontiers in Human Neuroscience doi:10.3389/fnhum.2015.00695.

Jagannathan, S.R., Ezquerro-Nassar, A., Jachs, B., Pustovaya, O.V., Bareham, C.A., Bekinschtein, T.A., 2018. Tracking wakefulness as it fades: Micro-measures of alertness. NeuroImage 176, 138–151. URL: https://doi.org/10.1016/j.neuroimage.2018.04.046, doi:10.1016/j.neuroimage.2018.04.046.

Kahana, M.J., 2006. The cognitive correlates of human brain oscillations. The Journal of neuroscience : the official journal of the Society for Neuroscience 26, 1669–1672. doi:10.1523/JNEUROSCI.3737-05c.2006.

Kamzanova, A.T., Kustubayeva, A.M., Matthews, G., 2014. Use of EEG workload indices for diagnostic monitoring of vigilance decrement. Human Factors doi:10.1177/0018720814526617.

Keitel, C., Thut, G., & Gross, J. (2017). Visual cortex responses reflect temporal structure of continuous quasi-rhythmic sensory stimulation. NeuroImage, 146(November), 58–70. doi: 10.1016/j.neuroimage.2016.11.043.

Kinchla, R.A., Wolfe, J.M., 1979. The order of visual processing: “Top-down,” “bottom-up,” or “middle-out”. Perception Psychophysics 25, 225–231. doi:10.3758/BF03202991.

Klimesch, W., Doppelmayr, M., Schimke, H., Pachinger, T., 1996. Alpha frequency, reaction time, and the speed of processing information. Journal of Clinical Neurophysiology doi:10.1097/00004691-199611000-00006.

Klug, M., Gramann, K., 2021. Identifying key factors for improving ica-based decomposition of eeg data in mobile and stationary experiments. European Journal of Neuroscience 54, 8406–8420.

Klug, M., Berg, T., & Gramann, K. (2024). Optimizing EEG ICA decomposition with data cleaning in stationary and mobile experiments. Scientific Reports, 14(1), 1–15. doi: 10.1038/s41598-024-64919-3

Kothe, C., Shirazi, S.Y., Stenner, T., Medine, D., Boulay, C., Grivich, M.I., Mullen, T., Delorme, A., Makeig, S., 2024. The Lab Streaming Layer for Synchronized Multimodal Recording. bioRxiv, 2024.02.13.580071URL: http://biorxiv.org/content/early/2024/02/14/2024.02.13.580071.abstract, doi:10.1101/2024.02.13.580071.

Ladouce, S., Darmet, L., Torre Tresols, J.J., Velut, S., Ferraro, G., Dehais, F., 2022. Improving user experience of ssvep bci through low amplitude depth and high frequency stimuli design. Scientific Reports 12, 8865.

Ladouce, S., Dehais, F., 2024. Frequency tagging of spatial attention using periliminal flickers. Imaging neuroscience 2, 1–17.

Ladouce, S., Tresols, J.T., Darmet, L., Ferraro, G., Dehais, F., 2021. Improving user experience of ssvep-bci through reduction of stimuli amplitude depth, in: 2021 IEEE International Conference on Systems, Man, and Cybernetics (SMC), IEEE. pp. 2936–2941.

Lamti, H.A., Khelifa, M.M.B., Alimi, A.M., Gorce, P., 2014. Influence of mental fatigue on p300 and ssvep during virtual wheelchair navigation, in: 2014 36th Annual International Conference of the IEEE Engineering in Medicine and Biology Society, IEEE. pp. 1255–1258.

Laukka, S.J., Järvilehto, T., I. Alexandrov, Y., Lindqvist, J., 1995. Frontal midline theta related to learning in a simulated driving task. Biological Psychology doi:10.1016/0301-0511(95)05122-Q.

Lee, K.A., Hicks, G., Nino-Murcia, G., 1991. Validity and reliability of a scale to assess fatigue. Psychiatry research 36, 291–298. doi:10.1016/0165-1781(91)90027-m.

Lim, J., Kwok, K., 2016. The Effects of Varying Break Length on Attention and Time on Task. Human Factors doi:10.1177/0018720815617395.

Lin, C.T., Chuang, C.H., Huang, C.S., Tsai, S.F., Lu, S.W., Chen, Y.H., Ko, L.W., 2014. Wireless and wearable EEG system for evaluating driver vigilance. IEEE Transactions on Biomedical Circuits and Systems 8, 165–176. doi:10.1109/TBCAS.2014.2316224.

Mackworth, N.H., 1948. The breakdown of vigilance during prolonged visual search. Quarterly journal of experimental psychology 1, 6–21.

Makeig, S., Bell, A., Jung, T.P., Sejnowski, T.J., 1995. Independent component analysis of electroencephalographic data. Advances in neural information processing systems 8.

Makri, D., Farmaki, C., Sakkalis, V., 2015. Visual fatigue effects on steady state visual evoked potential-based brain computer interfaces, in: 2015 7th International IEEE/EMBS Conference on Neural Engineering (NER), pp. 70–73. doi:10.1109/NER.2015.7146562.

Manly, T., Robertson, I.H., Galloway, M., Hawkins, K., 1999. The absent mind:: further investigations of sustained attention to response. Neuropsychologia 37, 661–670.

Martel, A., Dane, S., Blankertz, B., 2014. EEG predictors of covert vigilant attention. Journal of Neural Engineering doi:10.1088/1741-2560/11/3/035009.

Mehrabi, E., Kim, J.E., 2022. Physiological Measurements of Vigilance: A Systematic Review. Proceedings of the Human Factors and Ergonomics Society Annual Meeting doi:10.1177/1071181322661512.

Mehta, R.K., Parasuraman, R., 2013. Neuroergonomics: a review of applications to physical and cognitive work. Frontiers in human neuroscience 7, 889.

Möckel, T., Beste, C., Wascher, E., 2015. The Effects of Time on Task in Response Selection - An ERP Study of Mental Fatigue. Scientific Reports doi:10.1038/srep10113.

Molina, E., Sanabria, D., Jung, T.P., Correa, Á., 2019. Electroencephalographic and peripheral temperature dynamics during a prolonged psychomotor vigilance task. Accident Analysis and Prevention doi:10.1016/j.aap.2017.10.014.

Morgan, S. T., Hansen, J. C., & Hillyard, S. A. (1996). Selective attention to stimulus location modulates the steady-state visual evoked potential. Proceedings of the National Academy of Sciences of the United States of America. 10.1073/pnas.93.10.4770

Mouli, S., Palaniappan, R., 2016. Eliciting higher ssvep response from led visual stimulus with varying luminosity levels, in: 2016 International Conference for Students on Applied Engineering (ICSAE), pp. 201–206. doi:10.1109/ICSAE.2016.7810188.

Mylona, I., Glynatsis, M.N., Dermenoudi, M., Glynatsis, N.M., Floros, G.D., 2022. Validation of the Digital Eye Strain Questionnaire and pilot application to online gaming addicts. European journal of ophthalmology 32, 2695–2701. doi:10.1177/11206721211073262.

Müller, M. M., Picton, T. W., Valdes-Sosa, P., Riera, J., Teder-Sälejärvi, W. A., & Hillyard, S. A. (1998). Effects of spatial selective attention on the steady-state visual evoked potential in the 20–28 Hz range. Brain Research. Cognitive Brain Research, 6(4), 249–261. 10.1016/s0926-6410(97)00036-0.

Norcia, A.M., Appelbaum, L.G., Ales, J.M., Cottereau, B.R., Rossion, B., 2015. The steady-state visual evoked potential in vision research: A review. Journal of vision 15, 4–4.

O’Connell, R.G., Dockree, P.M., Robertson, I.H., Bellgrove, M.A., Foxe, J.J., Kelly, S.P., 2009. Uncovering the neural signature of lapsing attention: Electrophysiological signals predict errors up to 20 s before they occur. Journal of Neuroscience doi:10.1523/JNEUROSCI.5967-08.2009.

Oken, B.S., Salinsky, M.C., Elsas, S.M., 2006.Vigilance, alertness, or sustained attention: physiological basis and measurement. doi:10.1016/j.clinph.2006.01.017.

Onton, J., Delorme, A., Makeig, S., 2005. Frontal midline EEG dynamics during working memory. NeuroImage doi:10.1016/j.neuroimage.2005.04.014.

Ortner, R., Allison, B.Z., Korisek, G., Gaggl, H., Pfurtscheller, G., 2011. An SSVEP BCI to control a hand orthosis for persons with tetraplegia. IEEE Transactions on Neural Systems and Rehabilitation Engineering 19, pp. 1–5. doi:10.1109/TNSRE.2010.2076364.

Pan, Y., Frisson, S., Jensen, O., 2021. Neural evidence for lexical parafoveal processing. Nature Communications 12, 1–9.

Papadelis, C., Chen, Z., Kourtidou-Papadeli, C., Bamidis, P.D., Chouvarda, I., Bekiaris, E., Maglaveras, N., 2007. Monitoring sleepiness with on-board electrophysiological recordings for preventing sleep-deprived traffic accidents. Clinical Neurophysiology doi:10.1016/j.clinph.2007.04.031.

Parra, L., Sajda, P., 2003. Blind Source Separation via Generalized Eigenvalue Decomposition. The Journal of Machine Learning Research 4, 1261–1269.

Patterson Gentile, C., Aguirre, G.K., 2020. A neural correlate of visual discomfort from flicker. Journal of vision 20, pp. 1–10. doi:10.1167/jov.20.7.11.

Pershin, I., Candrian, G., Münger, M., Baschera, G.M., Rostami, M., Eich, D., Müller, A., 2023. Vigilance described by the time-on-task effect in EEG activity during a cued Go/NoGo task. International Journal of Psychophysiology doi:10.1016/j.ijpsycho.2022.11.015.

Peterson, D.J., Gurariy, G., Dimotsantos, G.G., Arciniega, H., Berryhill, M.E., Caplovitz, G.P., 2014. The steady-state visual evoked potential reveals neural correlates of the items encoded into visual working memory. Neuropsychologia 63, 145–153.

Pion-Tonachini, L., Kreutz-Delgado, K., Makeig, S., 2019. Iclabel: An automated electroencephalographic independent component classifier, dataset, and website. NeuroImage 198, 181–197.

Robertson, I.H., Manly, T., Andrade, J., Baddeley, B.T., Yiend, J., 1997. Oops!’: performance correlates of everyday attentional failures in traumatic brain injured and normal subjects. Neuropsychologia 35, 747–758.

Roy, R.N., Bonnet, S., Charbonnier, S., Campagne, A., 2013. Mental fatigue and working memory load estimation: interaction and implications for eeg-based passive bci, in: 2013 35th annual international conference of the IEEE Engineering in Medicine and Biology Society (EMBC), IEEE. pp. 6607–6610.

Roy, R.N., Charbonnier, S., Campagne, A., Bonnet, S., 2016. Efficient mental workload estimation using task-independent eeg features. Journal of neural engineering 13, 026019.

Russo, F. Di, & Spinelli, D. (1999). Spatial attention has different effects on the magno- and parvocellular pathways. NeuroReport, 10(13).

Shou, G., Dasari, D., Ding, L., 2015. Pre-stimulus alpha and post-stimulus N2 foreshadow imminent errors in a single task. Neuropsychologia 77, 346–358. URL: https://www.sciencedirect.com/science/article/pii/S0028393215301524, doi: 10.1016/j.neuropsychologia.2015.09.006.

Silberstein, R.B., Line, P., Pipingas, A., Copolov, D., Harris, P., 2000. Steady-state visually evoked potential topography during the continuous performance task in normal controls and schizophrenia. Clinical Neurophysiology 111, 850–857. doi:10.1016/S1388-2457(99)00324-7.

Silberstein, R.B., Schier, M.A., Pipingas, A., Ciorciari, J., Wood, S.R., Simpson, D.G., 1990. Steady-state visually evoked potential topography associated with a visual vigilance task. Brain topography 3, 337–347.

Small, A.J., Wiggins, M.W., Loveday, T., 2014. Cue-based processing capacity, cognitive load and the completion of simulated short-duration vigilance tasks in power transmission control. Applied Cognitive Psychology doi:10.1002/acp.3016.

Strijkstra, A.M., Beersma, D.G., Drayer, B., Halbesma, N., Daan, S., 2003. Subjective sleepiness correlates negatively with global alpha (8-12 Hz) and positively with central frontal theta (4-8 Hz) frequencies in the human resting awake electroencephalogram. Neuroscience Letters doi:10.1016/S0304-3940(03)00033-8.

Teng, F., Chen, Y., Choong, A.M., Gustafson, S., Reichley, C., Lawhead, P., Waddell, D., 2011. Square or Sine : Finding a Waveform with High Success Rate of Eliciting SSVEP Square or Sine : Finding a Waveform with High Success Rate of Eliciting SSVEP doi:10.1155/2011/364385.

Thomson, D.R., Besner, D., Smilek, D., 2015. A Resource-Control Account of Sustained Attention: Evidence From Mind-Wandering and Vigilance Paradigms. Perspectives on Psychological Science doi:10.1177/1745691614556681.

Tian, S., Wang, Y., Dong, G., Pei, W., Chen, H., 2018. Mental Fatigue Estimation Using EEG in a Vigilance Task and Resting States. Proceedings of the Annual International Conference of the IEEE Engineering in Medicine and Biology Society, EMBS 2018-July, 1980–1983. doi:10.1109/EMBC.2018.8512666.

Torkamani-Azar, M., Kanik, S.D., Aydin, S., Cetin, M., 2020. Prediction of Reaction Time and Vigilance Variability from Spatio-Spectral Features of Resting-State EEG in a Long Sustained Attention Task. IEEE Journal of Biomedical and Health Informatics 24, 2550–2558. doi:10.1109/JBHI.2020.2980056, arXiv:1910.10076.

Tran, Y., Craig, A., Craig, R., Chai, R., Nguyen, H., 2020. The influence of mental fatigue on brain activity: Evidence from a systematic review with meta-analyses. doi:10.1111/psyp.13554.

Trejo, L.J., Kubitz, K., Rosipal, R., Kochavi, R.L., Montgomery, L.D., 2015. EEG-Based Estimation and Classification of Mental Fatigue. Psychology doi:10.4236/psych.2015.65055.

Wang, C., Trongnetrpunya, A., Samuel, I.B.H., Ding, M., Kluger, B.M., 2016. Compensatory neural activity in response to cognitive fatigue. Journal of neuroscience 36, 3919–3924.

Ward, A.F., Wegner, D.M., 2013. Mind-blanking: When the mind goes away. Frontiers in Psychology 4, 1–15. doi:10.3389/fpsyg.2013.00650.

Warm, J.S., Parasuraman, R., Matthews, G., 2008. Vigilance requires hard mental work and is stressful. doi:10.1518/001872008×312152.

Wascher, E., Rasch, B., Sanger, J., Hoffmann, S., Schneider, D., Rinkenauer, G., Heuer, H., Gutberlet, I., 2014. Frontal theta activity reflects distinct aspects of mental fatigue. Biological Psychology 96. doi:10.1016/j.biopsycho.2013.11.010.

Wu, C.H., Lakany, H., 2013. The Effect of the Viewing Distance of Stimulus on SSVEP Response for Use in Brain-Computer Interfaces, in: 2013 IEEE International Conference on Systems, Man, and Cybernetics, IEEE. pp. 1840–1845. doi:10.1109/SMC.2013.317.

Xie, J., Xu, G., Wang, J., Li, M., Han, C., Jia, Y., 2016. Effects of mental load and fatigue on steady-state evoked potential based brain computer interface tasks: a comparison of periodic flickering and motion-reversal based visual attention. PloS one 11, e0163426.

Xie, T., Li, M., Hao, C., Peng, Y., Luo, W., Ma, N., 2023. How the time-of-day affects the EEG signatures of vigilance fluctuation. Chronobiology International 40, 1059–1071. URL: https://doi.org/10.1080/07420528.2023.2250439, doi:10.1080/07420528.2023.2250439.

Zemon, V., Gordon, J., 2006. Luminance-contrast mechanisms in humans: visual evoked potentials and a nonlinear model. Vision research 46, pp. 4163–4180. doi:10.1016/j.visres.2006.07.007.

Zheng, B., Tien, G., Atkins, S.M., Swindells, C., Tanin, H., Meneghetti, A., Qayumi, K.A., Neely, O., Panton, M., 2011. Surgeon’s vigilance in the operating room. American journal of surgery 201, 673–677. doi:10.1016/j.amjsurg.2011.01.016.

Zhigalov, A., Herring, J.D., Herpers, J., Bergmann, T.O., Jensen, O., 2019. Probing cortical excitability using rapid frequency tagging. NeuroImage 195, 59–66.

Zhu, D., Bieger, J., Molina, G.G., Aarts, R.M., 2010. A Survey of Stimulation Methods Used in SSVEP-based BCIs. Computational Intelligence and Neuroscience 2010, pp. 1–12. doi:10.1155/2010/702357.

